# Network Dynamics and State-Dependent Effects of Electrical Stimulation in Recurrent Excitatory-Inhibitory Populations

**DOI:** 10.64898/2026.07.29.741519

**Authors:** Spandan Sengupta, Shervin Safavi, Thomas R. Knösche, Milad Lankarany

## Abstract

Deep Brain Stimulation (DBS) is an established clinical treatment for a variety of neurological disorders, including Parkinson’s Disease where it has been shown to reduce motor symptoms as well as disrupt pathological beta oscillations in the basal ganglia. The mechanisms of action of DBS on the collective activity of neuronal circuits is not fully understood. We use a recurrently-connected excitatory-inhbitory network based on the Brunel network architecture that can produce activity in a variety of states. Using a model of DBS that can reproduce observed effects such as antidromic activation, local somatic suppression, and axonal activation, we characterize the effect of stimulation across the entire parameter space of the network. We show that the effects of stimulation are dependent on the baseline state of the network, with the level of beta suppression dependent on the level of inhibition and the external drive. Specifically, networks with higher inhibition and lower drive show greater disruption of beta oscillations. We further show that networks in different states are preferentially sensitive to different frequencies of stimulation, suggesting that alternative protocols to the clinically standard high-frequency stimulation may have therapeutic efficacy.

## 1 Introduction

Deep brain stimulation (DBS) has emerged as a highly successful therapeutic intervention for the treatment of several neurological disorders, including Parkinson’s disease, essential tremor, dystonia, and Alzheimer’s disease[1–5]. The clinical efficacy of high-frequency DBS in mitigating symptoms of Parkinson’s disease is correlated with the suppression of pathological, hyper-synchronous oscillations within the basal ganglia, particularly in the beta band (13-30 Hz)[3, 6–8]. However, despite its widespread clinical usage, a comprehensive mechanistic understanding of how the externally applied stimulation interacts with the endogenous network states remains elusive. While a significant amount of experimental and computational work has gone into studying the effect of DBS at the local level[9–14], the effect on the network’s collective dynamics is relatively unexplored. An understanding of these DBS-modulated network dynamics could allow clinical protocols to be more specific based on the underlying pathology. Variability in the clinical outcome of the DBS intervention suggests that an understanding of the relationship between the dynamical regime of the network and the functional efficacy of the intervention is crucial for more consistent positive clinical outcomes.

A major hurdle in mapping stimulation-network interactions is the lack of robust models, specifically, network models that can replicate diverse activity states and stimulation models that can account for the paradoxical effects of DBS. While some models consider the impact of DBS by injection of a supra-threshold current[10, 15], they can fail to capture important observed effects such as antidromic activation [16] and somatic-axonal decoupling[9]. We use a model of stimulation that involves activation of synapses, producing stimulation-induced action potentials in efferent and afferent neurons of the stimulated population. Using this model implemented in a modified Brunel network architecture[17], we are able to systematically characterize the effects of stimulation across the entire two-dimensional parameter space of relative inhibitory weight and external drive. We explore the effects of stimulation on the collective dynamics of the network such as population firing rates, firing regularity, population synchrony, and beta-band spectral power, across the diverse regimes of activity the network can exist in. We evaluate the network’s response to external stimulation across a range of stimulation frequencies and find state-specific differences in the response to high- and low-frequency stimulation.

Our results suggest that the response to external stimulation depends not only on the observable activity of the network, but also on the underlying properties of its structure and connectivity. We demonstrate that networks with threshold and sub-threshold levels of drive show the strongest presence of beta-band activity, and are also the most sensitive to stimulation at lower frequencies. Conversely, highly driven, inhibition-dominant networks require higher frequencies of stimulation to show functional improvements. Our findings offer a framework for predicting the efficacy of DBS protocols with variable intrinsic activity, providing a mechanistic foundation for further exploring the effects of different stimulation protocols based on the baseline state.

## 2 Results

We systematically investigated the effect of electrical stimulation on the Brunel Network. We first characterize the baseline dynamics of the Brunel Network when no stimulation is applied. We then introduce the necessary modifications that are required to implement the model of stimulation, and the consequent effects on the network’s dynamics without the application of stimulation. We use four metrics that capture the firing activity of individual neurons as well as the global activity of the entire population: Population Firing Rate, Coefficient of Variation (CV) of Inter-spike Intervals (ISIs), Population Synchrony (as pair-wise cross-correlation of spike trains), and the power in the Beta frequency band. We then show how high-frequency electrical stimulation at the clinical standard of 130 Hz [18] affects the activity of individual neurons as well as the collective dynamics of the population at different configurations of the network. We proceed to do an extensive parameter sweep across a wide range of relative inhibitory strength *g* and relative external drive *ν* when the network is stimulated at high, medium, and low frequency. We show the effects of different frequencies of stimulation on the aforementioned metrics, with an emphasis on the capacity of stimulation to suppress beta-band activity. We show how the stimulation frequency required to produce changes in the collective dynamics of the network is dependent on the intrinsic properties of the network (Figure 1).

**Figure 1:**
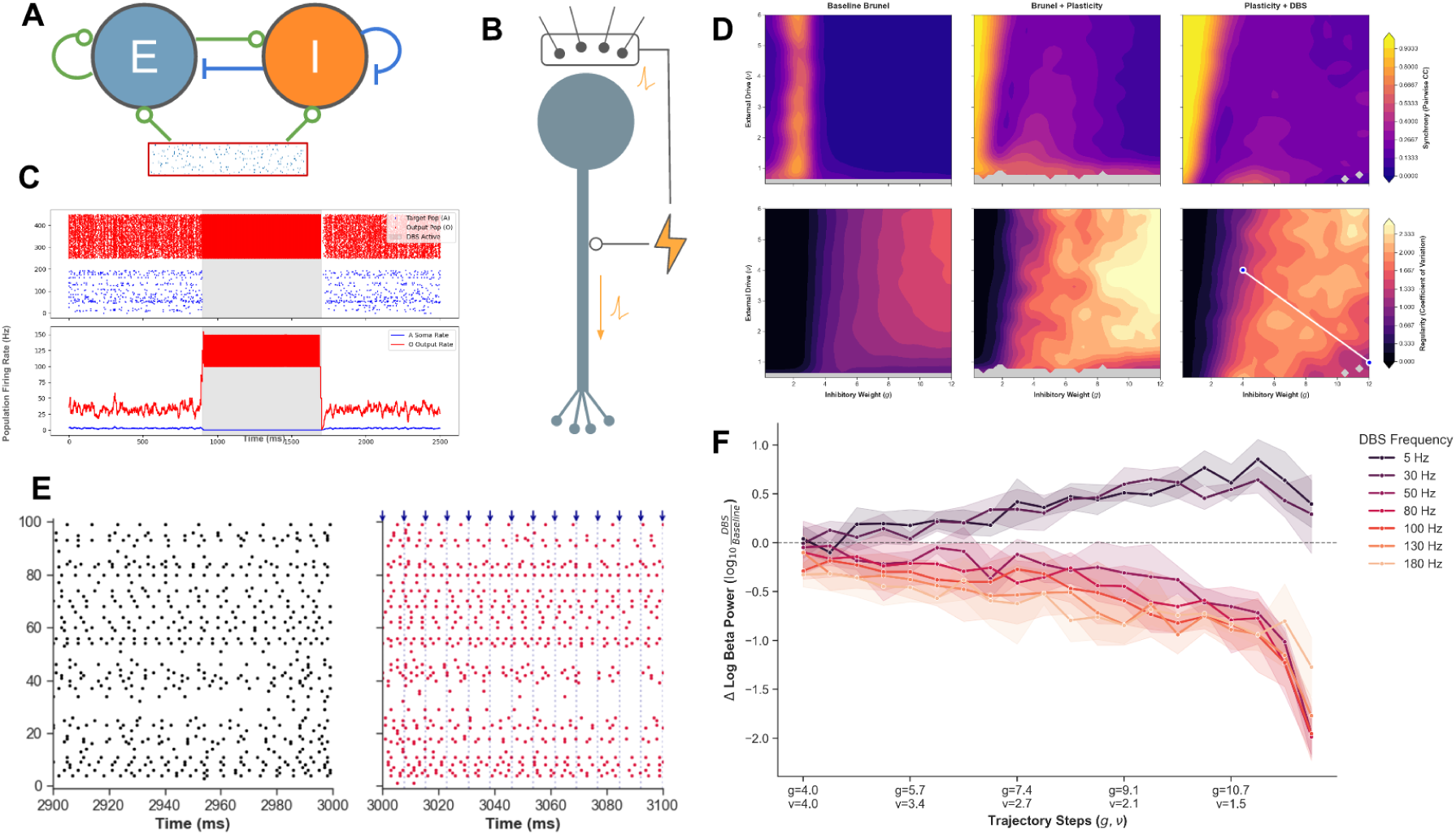
A: Schematic of the network architecture – an excitatory (E) and inhibitory (I) population with sparse connections to each other and themselves, receiving common external input from a Poisson spike train. A subset of the excitatory population is the target of stimulation. B: Schematic of the stimulation model – each stimulation pulse activates the presynaptic inputs of the target as well as its downstream targets. C: Spike Raster (top) and Population Firing Rate (bottom) of a population with dominant inhibitory afferents that is the target of stimulation, and a downstream population (red) receiving excitatory inputs from the target population. Stimulation causes suppression of firing in the target population while increasing activity in the downstream population. D: Comparison of the Brunel network, the plastic Brunel network with short-term depression, and the plastic Brunel network under the effect of Deep Brain Stimulation. The variation in Population Synchrony (top) and CV of ISIs (bottom) across the parameter space of *g* and *ν*. E: Spike Raster of a network at Baseline (left) and under the effect of DBS (right). Pulse timings are shown as blue arrows. F: Change in the Reduction of Beta-band Power along a Trajectory in the parameter space (marked in bottom right plot in D) at different frequencies of stimulation.

### 2.1 The Brunel Network

We provide a short summary of the observed characteristics of the Brunel network, specifically the variation of the mean firing rate and the coefficient of variation (CV) of inter-spike intervals (ISIs) with the control parameters of the network: the relative level of inhibition *g*, and the relative rate of the external Poisson input *ν*.

The relative strength of inhibition is the primary driver of the network’s phase transitions. The *g <* 3, the network remains in an excitation-dominated state where the firing rate is high and neurons fire close to their maximal firing potential. As *g* decreases, the recurrent inhibition begins to reduce the firing rate, leading to a sharp non-linear decrease in the firing rate. In the excitation-dominated regime, the CV of the ISIs is near zero, indicating highly regular and periodic firing. The *g* increases, and the network transitions into the inhibition-dominated state, the CV increases sharply. This corresponds to the spike timings becoming more irregular. At higher values of the external drive, the CV continues to increase after the phase transition with increasing *g*. The network activity shows the emergence of irregular population bursts, where a large number of neurons fire in phase with one another. The external drive has an impact on both the firing rate and the CV of ISIs in the inhibition-dominant state. An increase in *ν* causes a linear increase in the firing rate of the population, with the effect being stronger for more excitatory states. The impact on the regularity of firing is also dependent on the level of inhibition in the network. More excitatory networks, show a decrease in CV with increasing *ν*, with the activity becoming close to a Poisson process with *CV* = 1. More inhibitory networks show an increase in CV with increasing *ν* with the emergence of irregular population bursts.

### 2.2 Addition of Exponential Synapses and Short-term Plasticity

Exponential synapses introduces a synaptic time constant, meaning a post-synaptic current decays over time instead of instantaneously, as is the case in the original Brunel network. This causes some changes in the activity of the network as its parameters are varied. Although the overall dynamics of the network remain consistent with the original Brunel network, the scales of some of its features change. In the excitation-dominant regime (*g <* 3), the firing rates are significantly higher, as the sustained excitatory currents allow the network to maintain faster firing. In the inhibition-dominant state, the variation of the CV of ISIs with increasing *g* is more dramatic. At high drive, the CV goes up drastically with increased inhibition, with the network exhibiting stronger bursting activity. At low drive, the CV falls sharply as well, reaching values close to zero. The strong, persistent inhibition causes the population to be close to silence, with regular coordinated bursts of activity (Figure 2).

**Figure 2:**
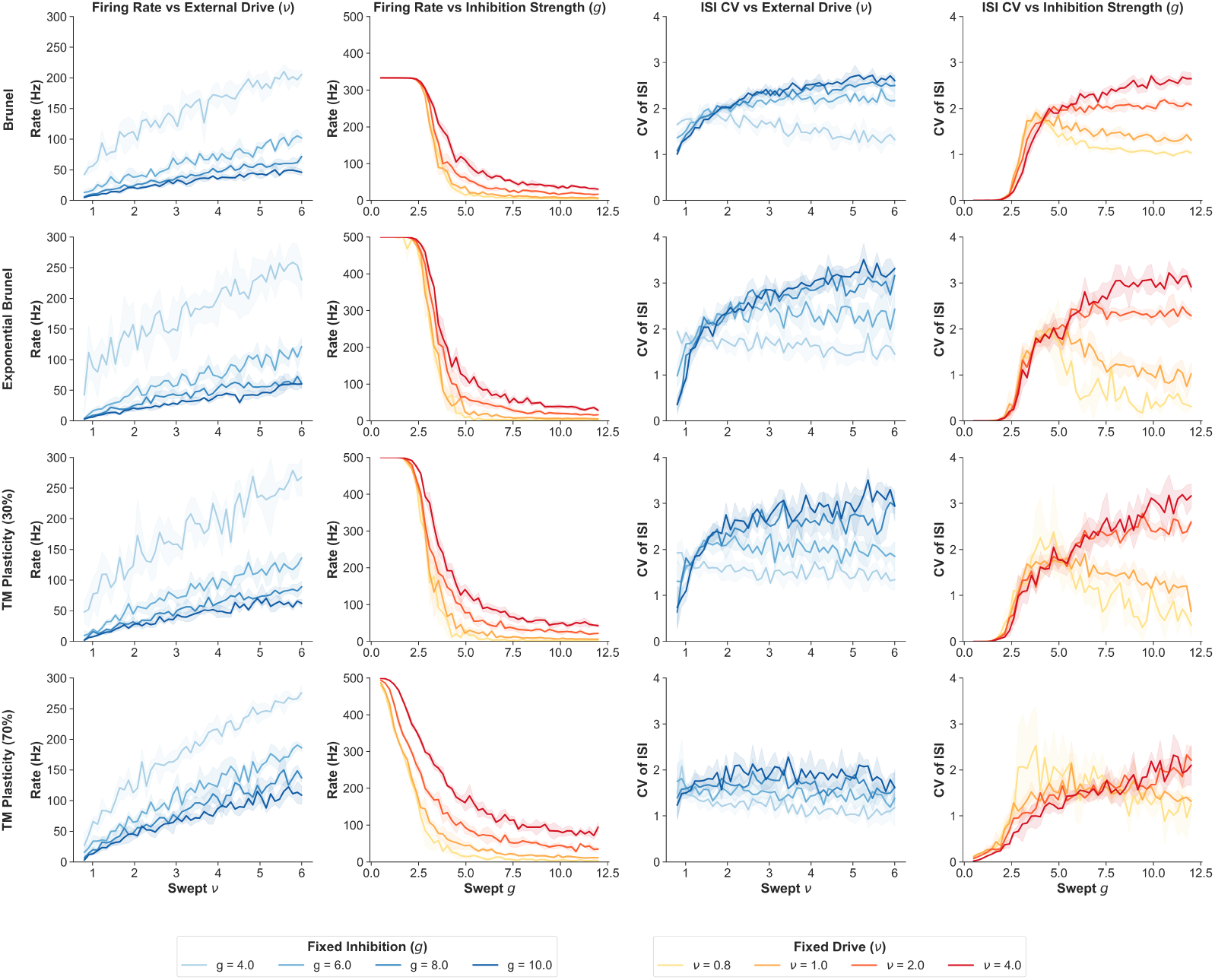
The variation of the Mean Population Firing Rate and Coefficient of Variation (CV) of Inter-Spike Intervals (ISIs) with the control parameters of the network: relative weight of inhibitory connections (*g*) and relative rate of external Poisson input (*ν*) for the Brunel network (top row), the Brunel network with Exponential Synapses (2nd row), the Brunel network with 30% Short-term Depressing Tsodyks-Markram (TM) synapses (3rd row), and the Brunel network with 70% Short-term Depressing TM synapses.

We also observed distinct network dynamics depending on the level of synaptic depression. At low levels of short-term depression: when 30% of the synapses in the network are plastic, the network’s response to change in parameters is very similar to when all synapses are exponential. The major change observed is the increased variability between trials, as the network becomes more sensitive to initial conditions. At high levels of synaptic depression, when 70% of the synapses are plastic, the network’s dynamics shift considerably. One important distinction is the transition from the excitation-dominant state to the inhibition-dominant state that is characterised by a sharp decrease in firing rate and increase in CV of ISIs. With the addition of synaptic plasticity, these transitions are less sharp and occur over a larger range of *g* values. In the inhibition-dominant state, the CV of ISIs is more homogenous: we do not observe the distinct trends for low and high levels of external drive as before. There is less bursting activity observed in the inhibition-dominant state, and it is characterised by asynchronous irregular activity across a large range of parameters. At highly inhibitory states, the firing rate remains higher than previously observed. The depression of inhibitory synapses reduces their capacity to suppress the firing activity, allowing for a higher firing rate. The increased short-term depression leads to a loss of distinct regimes in the inhibition-dominant state, as was previously characterised by the regularity of the firing activity, creating a network that is less sensitive to parameter changes.

#### 2.2.1 The Effects of Plasticity Across the 2D Parameter Space

Looking at the changes in the population dynamics of the network across the entire parameter space of *g* and *ν*, we focus on four metrics: Population Firing Rate, CV of ISIs, Population Synchrony (as pair-wise cross-correlation of spike trains), and the Power in the Beta Frequency Band. As we observed previously, there is a sharp transition between an excitation-dominant and inhibition-dominant state that occurs close to *g* = 3, characterised by a drastic change in firing rate and CV, as well as a sharp decrease in synchrony. The excitation-dominant state is characterised by high synchrony as the entire population fires in a highly-coordinated fast-firing regime. In the inhibition-dominant state, we observe that networks with high external drive and inhibition (in the top-right corner) show high CV, characterised by irregular population bursts. Networks with lower external drive show lower rates of firing with CV close to one, indicative of Poisson-like irregular asynchronous activity (Figure 3).

**Figure 3:**
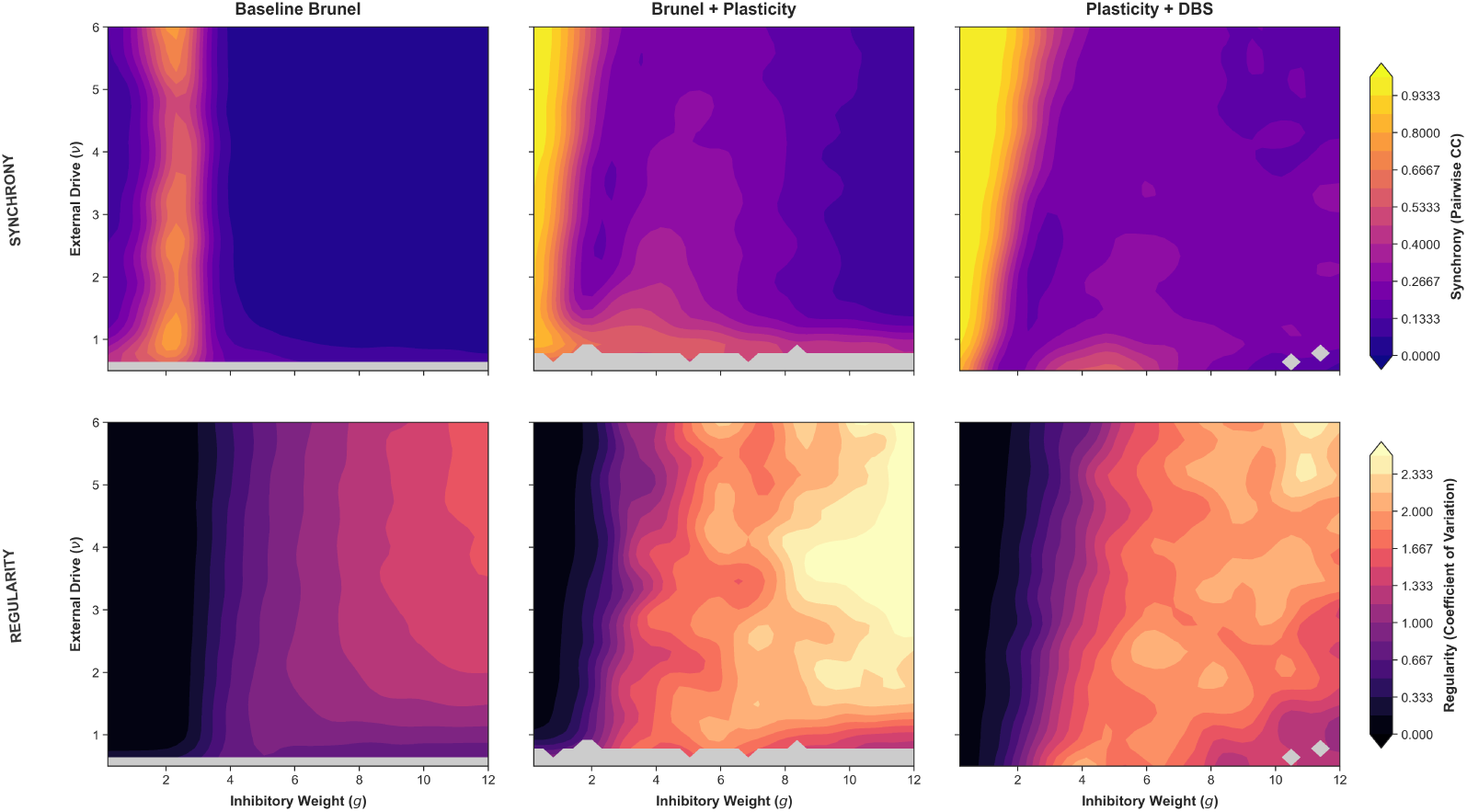
The variation in the Population Synchrony and the CV of ISIs for the Brunel network, the plastic Brunel network with short-term depression, and the plastic Brunel network under the effect of Deep Brain Stimulation. Top panels illustrate topological changes in global pairwise synchrony, while bottom panels display corresponding changes in spike-timing regularity (CV).

The introduction of short-term plasticity and exponential synapses shows a significant breakdown in phase boundaries observed in the original Brunel network. The exhaustion of synaptic resources in response to increased firing activity acts as a means of regularizing population bursting dynamics and homogenizing the activity in the inhibition-dominant regime. The boundary of the strongly synchronous fast-firing state is shifted to the left, especially for lower external drive. The transition from this state to asynchronous slower-firing is also more gradual. Immediately after this transition, there is a band of asynchrony, followed by a wedge of synchronous activity between *g* = 4 and *g* = 6. Additionally, at very low external drive, there is a band of strong synchronous activity with strong beta power and high irregularity. This activity state is characterised by strong population bursts from a large number of neurons that occur 50-80 ms apart. This coordinated population activity causes the presence of the abnormally strong power in the beta-frequency band. In the inhibition-dominant regime, there is a marked decrease of irregularity compared to the Brunel network. The strongly irregular activity observed in networks with high inhibition and high external drive with CV approaching 1.5 is no longer observed. In this plastic network, the highest irregularity occurs in networks with high inhibition and low external drive (bottom left of plot) with CV values close to 1. These networks are characterised by activity that is asynchronous and irregular with occasional bursts of activity from a small set of coordinated neurons.

### 2.3 High Frequency Stimulation Effects are Dependent on the Intrinsic Properties of the Network

We look at the effect of high-frequency electrical stimulation at 130 Hz on the firing patterns of the network as well as the Local Field Potential (LFP) to understand how the stimulation produces qualitative changes on the network’s dynamics. We try to characterize these changes in terms of their relevance to therapeutic efficacy through reduction of the beta power as well as changes in the synchrony and regularity of the firing (Figure 4).

**Figure 4:**
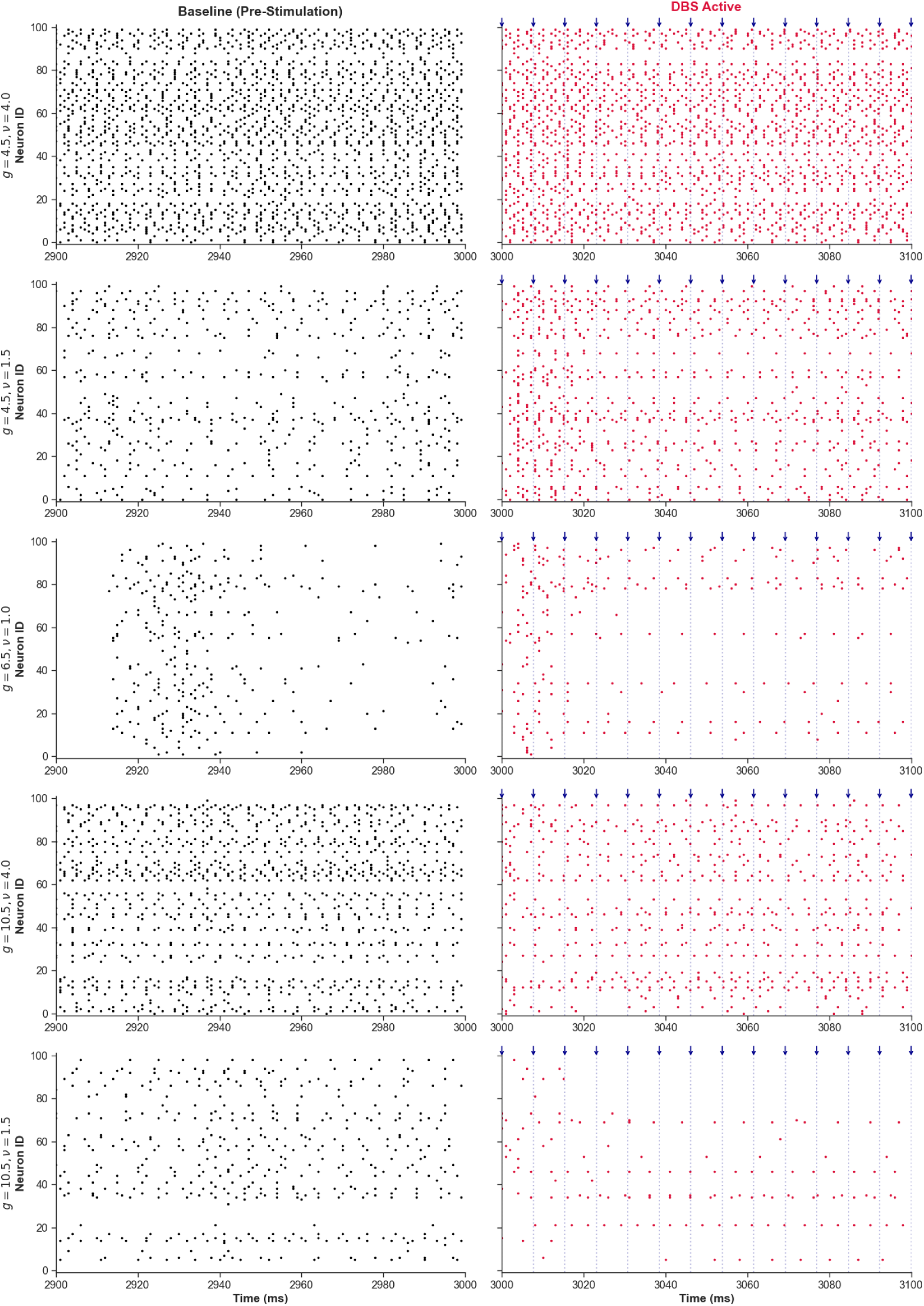
Change in firing activity under high-frequency DBS across representative parameter regimes. Spiking activity from a sub-sampled population of 100 neurons is shown over a 100 ms window before stimulation (Left, Baseline) and immediately after stimulation onset (Right, DBS Active) across five (*g, ν*) parameter coordinates. Dark blue arrows and vertical dashed lines denote the times of individual DBS pulses (130 Hz)

#### Balanced Inhibition, High Drive (*g* = 4.5*, ν* = 4.0): Entrainment and Increased Firing Irregularity

At *g* = 4.5, *ν* = 4.0, the application of high-frequency stimulation causes a transition from regular, bursting activity to a regime characterised by entrainment to the stimulation frequency. The Power Spectral Density (PSD) shows a sharp, narrow-band spike at 130 Hz, indicating strong network entrainment. In the beta-band, the power spectrum when stimulation is applied is lower than the baseline, suggesting that the stimulation is successful in disrupting the intrinsic slow oscillations. This is further evident when comparing the Local Field Potential (LFP) traces in the baseline and stimulated states, which goes from slow, low-amplitude oscillations to a high-frequency regular oscillation. However, the regularity of the individual neurons’ firing is decreased under the influence of stimulation, with the CV of ISIs increasing from 1.98 to 2.53, indicating that individual neurons are often skipping cycles of the stimulation-entrained oscillation. While the 130 Hz stimulation regularizes the network’s collective dynamics by suppressing low-frequency fluctuations and entraining it to high-frequency regime, it paradoxically makes the spiking of individual neurons more irregular.

#### Balanced Inhibition, Low Drive (*g* = 4.5*, ν* = 1.5): Regularization of Global and Local Activity

At *g* = 4.5, *ν* = 1.5, the baseline activity of the network is more sparse with irregular population bursts. This is shown in the CV of ISIs that has a higher value of 4.5. The introduction of stimulation to this network has a regularizing effect on the individual neurons’ firing as well as the collective dynamics of the network. While in the baseline states, some neurons have long ISIs around 12-15 ms. Under the effect of stimulation, the distribution of ISIs is narrowed and shifted to the left, with an increase in shorter ISIs. Similar to the balanced network with high drive, the stimulation is effective at entraining the network activity to the stimulation frequency, with a sharp peak in the PSD visible at 130 Hz. It also has the effect of suppressing low-frequency activity in the beta-band. In contrast to the network with higher drive, the stimulation regularizes both the collective activity of the network as well as the individual firing of the neurons.

#### High Inhibition, High Drive (*g* = 10.5*, ν* = 4.5): Suppression of Firing Rate and Increased Regularity

At *g* = 10.5 and *ν* = 4.5, the opposing forces of the external drive and inhibition makes the network behave similarly to the balanced network with low drive. It is characterised by sparse firing with irregular population bursts and a resultant high CV of ISIs. The effect of stimulation is similar to the balanced, low drive network as well: there is an observable reduction in low frequency oscillations, as well as entrainment at the stimulation frequency. However, due to the high inhibition in the network, the stimulation causes a significant reduction in firing rate, with a number of neurons being silent while the network is stimulated. The stimulation also regularizes the firing of the individual neurons, with a reduction of CV of ISIs to 3.09. When stimulation is applied to a irregularly bursting network, it has a regularizing effect. However, the specific balance of inhibition and external input can affect the change it produces in the firing rate.

#### High Inhibition, Low Drive (*g* = 10.5*, ν* = 1.5): Strong Suppression of Beta-band Activity and Suppression of Firing Rate

At *g* = 10.5 and *ν* = 1.5, the network exhibits extremely sparse, inhibition-dominated activity. At this configuration, the addition of high-frequency stimulation further silences the population, making the population activity almost exclusively driven by the stimulation. Under the influence of stimulation, several neurons are completely silent, due to the activation of the large number of inhibitory synapses. Some neurons firing regularly, driven by the stimulation. The population activity is characterized by strong entrainment to the 130 Hz stimulation, and almost complete suppression of any low frequency activity. Compared to the previous networks, this shows the greatest effect in reduction of beta-band power, but is also accompanied by significant reduction in firing rate.

### 2.4 Stimulation-Induced Effects Across the 2D Parameter Space

We further characterize the effect of stimulation across the entire 2-dimensional parameter space of *g* and *ν* to identify regions that respond differently to the applied stimulation. We first characterize the effect of high-frequency stimulation at 130 Hz, and subsequently compare the effects of lower frequencies such as 50 Hz and 5 Hz stimulation (Figure 5).

**Figure 5:**
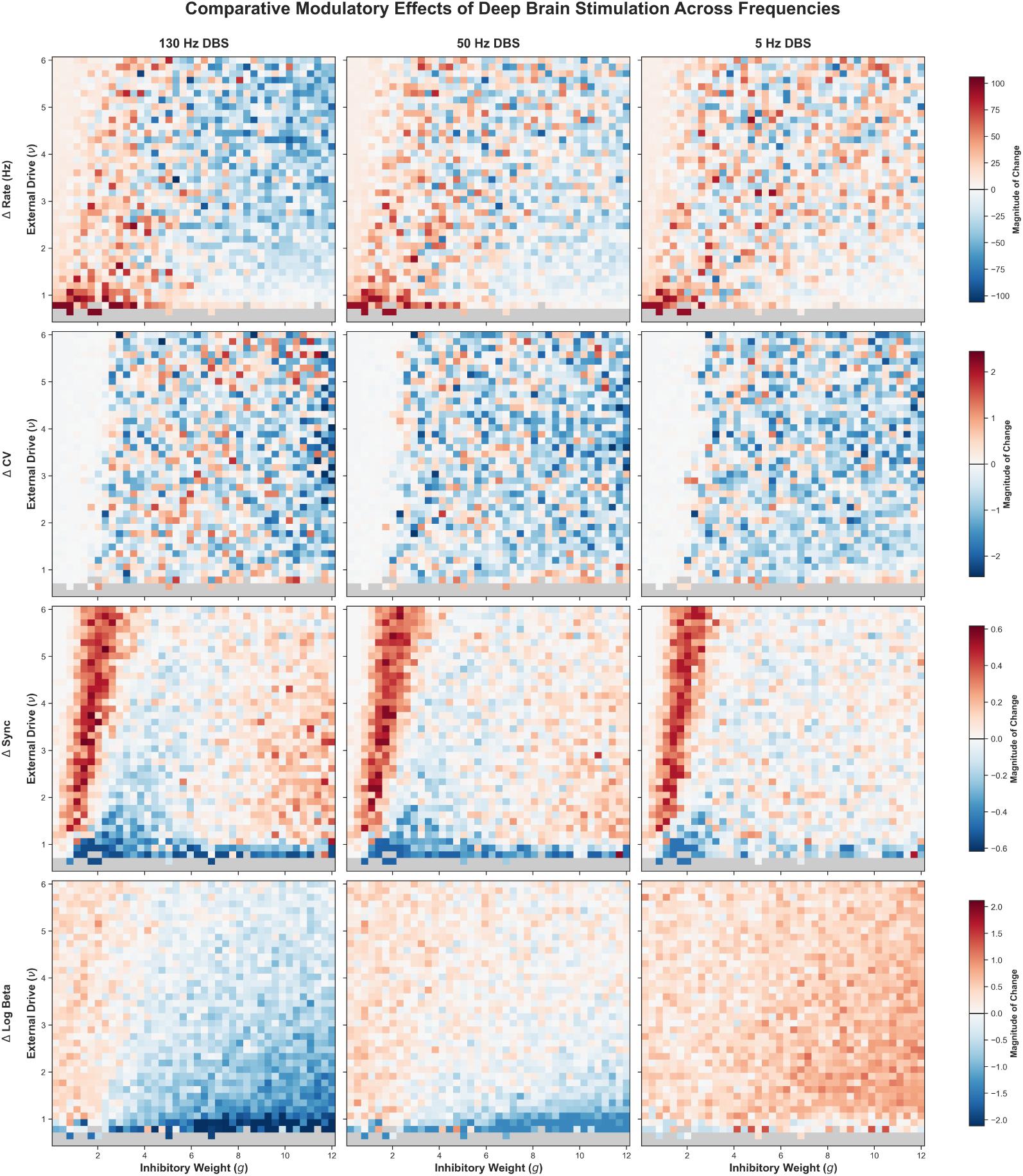
Frequency-dependent impacts of DBS across circuit activity metrics. Parameter sweeps across relative inhibitory strength (*g*) and external drive (*ν*) showing the change (Δ) in mean firing rate (row 1), coefficient of variation (ΔCV, row 2), pairwise synchrony (ΔSync, row 3), and log beta power (ΔLog Beta, row 4) under three distinct stimulation frequencies: 130Hz (left), 50 Hz (middle), and 5 Hz (right). High-frequency stimulation (130 Hz) driving into highly inhibited regimes (*g >* 4) demonstrates maximum therapeutic efficacy via widespread beta band suppression and desynchro-nization. In contrast, low-frequency stimulation (5 Hz) drives resonant amplification of beta power across the state space. Non-active or silent network states are masked in neutral gray.

#### Effect of High-Frequency Stimulation (130 Hz)

The effect of the high frequency stimulation on the mean population firing rate is largely dependent on the excitatory-inhibitory balance of the network. Networks that are dominated by excitation show an increased firing rate, while networks with dominant inhibition show decreased firing in response to 130 Hz stimulation. The external drive also has a secondary effect, with low drive states with high excitation showing the greatest increase in firing rate, and conversely, highly inhibitory states with a high level of external drive show the highest firing rate reduction. The transition between networks that show an increase in firing and those that show a decrease is gradual, with no sharp boundary observable in the 2D parameter space. The transition occurs close to the level of inhibition in a balanced network.

For networks dominated by a high level of excitation, the addition of 130 Hz stimulation does not have an effect on the regularity of firing. These networks are characterized by highly regular, fast, synchronous firing which persists even after the application of stimulation. Outside this regime, the effect of stimulation on the regularity of firing is mottled over the parameter space, with no regions of similar effect observable. However, networks with a higher level of inhibition are biased to be more likely to be regularized in response to the high-frequency stimulation.

Three major regions are identifiable when the change in synchrony in response to 130 Hz stimulation is mapped over the parameter space. The low inhibition regime, characterized by regular, fast activity is further synchronized by the stimulation. Across a wide range of *g*, networks with low external drive (*ν <*= 1) show strong desynchronization when high-frequency stimulation is applied. These networks intrinsically have highly synchronous population bursts that are disrupted by the addition of high-frequency stimulation. Networks with high inhibition (*g >* 8) also show an increase in synchrony when stimulated. This occurs for a wide range of external drives above the threshold level. In between the highly excitatory and highly inhibitory states that show an increase in synchrony, there is a wedge in the parameter space that shows decreased synchrony in response to high-frequency stimulation. This occurs at balanced levels of excitation and inhibition (2 *< g <* 6) and the effect is more pronounced at lower external drive.

Strongest reduction in beta power occurs for networks with low external drive. These states have intrinsically high beta due to the synchronous population bursts. These are disrupted by high-frequency stimulation leading to a large decrease in the beta power. The reduction of beta power is increases with increase in the level of inhibition and decrease in the external drive. The lower right corner shows the strongest beta power reduction which decreases as one moves away from it. For excitation dominated networks, the applied stimulation can lead to an increase in beta power. However, these networks have very low beta power intrinsically and the activity is dominated by high-frequency firing in both baseline and stimulated states.

#### Effect of Mid-Frequency Stimulation (50 Hz)

Under the effect of 50 Hz stimulation, the boundary between network states that show an increase and those that show a decrease in firing activity becomes less clear. A larger section of the parameter space shows an increase in firing in response to stimulation, most notably in networks with high inhibition and high external drive that show a reduction in firing rate under 130 Hz stimulation. The effect of 50 Hz stimulation on regularization of the firing activity is more consistent. While the highly excitatory networks are still unaffected by stimulation at 50 Hz, networks with *g >* 2 are more likely to be regularized by the stimulation. At this frequency, the stimulation is less effective at modulating the level of synchrony in the population. The effects on synchrony follow the same patterns as in the case of 130 Hz stimulation, but with the strength of the effect reduced. This reduced strength of effect also occurs in the case of beta-band power reduction. The 50 Hz stimulation, while having a strong effect on networks with very low external drive that have high beta power, fails to decrease beta power as efficiently over the rest of the parameter space. For networks with a high level of external drive, 50 Hz stimulation can have an amplifying effect of the beta band activity. While it does have a suppressing effect on networks with high inhibition and low external drive, the reduction in power is weaken than with applied 130 Hz stimulation.

#### Effect of Low-Frequency Stimulation (5 Hz)

At low frequency stimulation, the effect of the stimulation on the firing rate and regularity of the firing is similar to the case of 50 Hz stimulation. The same patterns of firing rate amplification for excitatory networks and networks with high external drive, and firing rate reduction for networks with high inhibition and low external drive are still observed. The regularity of firing is increased for a wide range of network parameters, with the highly excitatory networks unaffected as before. Similar changes to the synchrony of the network is also observed, with highly excitatory and highly inhibitory networks showing an increase in synchrony while balanced networks show a decrease in synchrony in response to stimulation. The range of *g* values that show decreased synchrony is slightly enlarged, with networks up to *g* = 8 being desynchronized by stimulation. The most notable difference observed is the effect of stimulation on the beta band activity. Contrary to the stimulation at 50 Hz and 130 Hz, 5 Hz stimulation shows no beta suppression across the entire parameter space. Instead, this frequency of stimulation amplifies the beta band activity across the whole range of *g* and *ν* values, with the strongest amplification occurring for networks with high *g* and high *ν*.

### 2.5 The Effect of Stimulation Frequency on Beta-band Suppression is Dependent on the External Drive

Across the entire parameter space, low-frequency stimulation uniformly exerts an amplifying effect on intrinsic beta-band (13–30 Hz) spectral power. This amplification is maximally pronounced when the stimulation frequency falls within the bounds of the beta band, driving a resonant entrainment mechanism phase-locked to the stimulation pulses. Highly excitatory, saturated networks display minimal change from external stimulation. In these regimes, characterized by low relative inhibition (*g <*= 2), the network’s intrinsic dynamics are dominated by highly synchronous, high-frequency, regular firing. This state shows negligible spectral or temporal alterations even when subjected to high-frequency stimulation (Figure 6).

**Figure 6:**
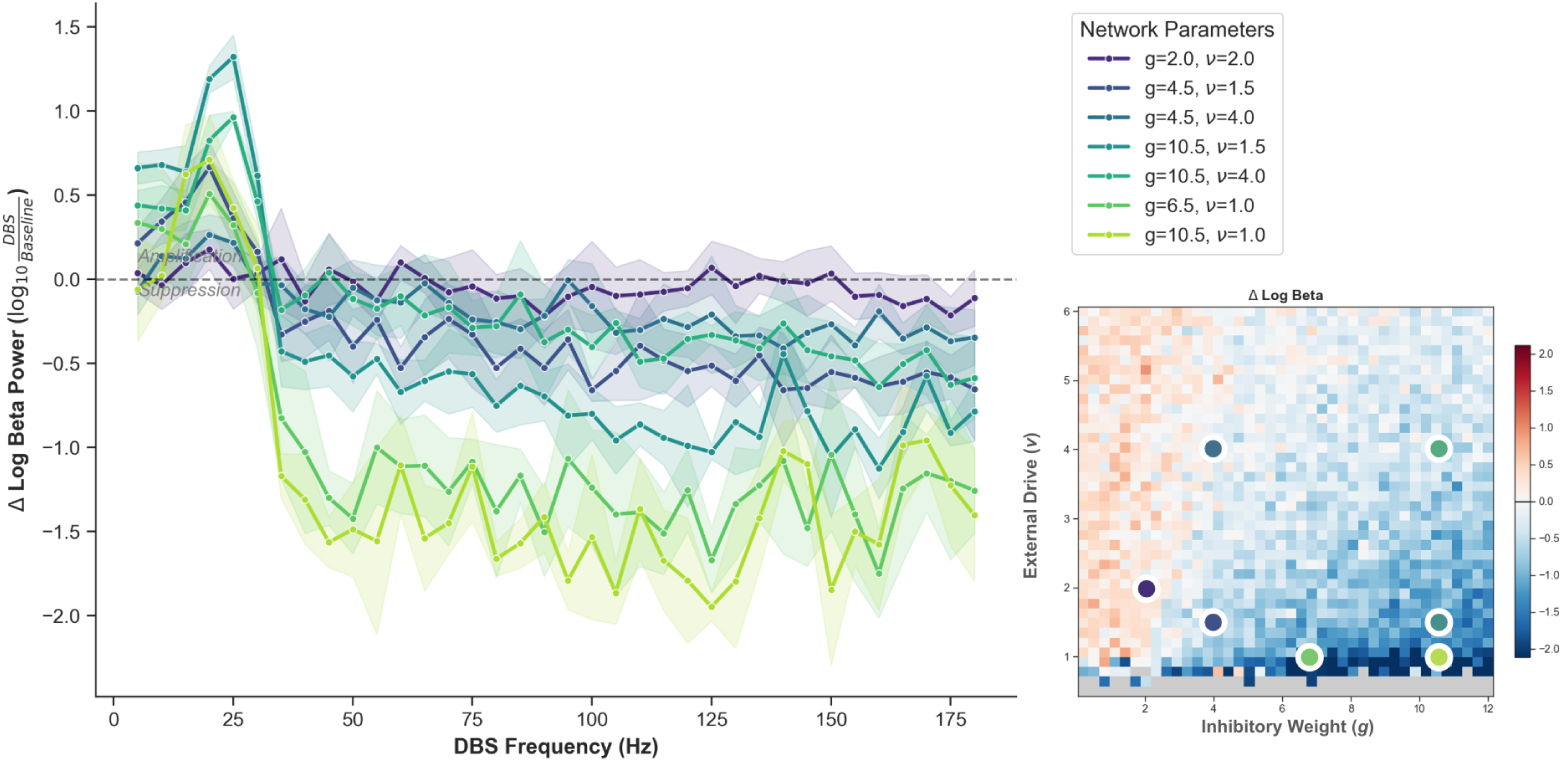
Frequency-dependent modulation of pathological beta oscillations via Deep Brain Stimulation (DBS) across diverse network regimes. (Left) Relative change in local field potential (LFP) log beta power (ΔLog Beta Power = log_10_ Beta_DBS_ *−* log_10_ Beta_Baseline_) plotted as a function of DBS stimulation frequency (5–180 Hz) for seven representative coordinates in the (*g, ν*) parameter space. Shaded regions denote *±*SEM across simulation trials. Low-frequency stimulation (*<* 35 Hz) induces prominent resonant amplification of beta-band activity across most responsive regimes, peaking near 25 Hz. Conversely, high-frequency stimulation (*>* 100 Hz) systematically suppresses beta power, with the strongest therapeutic efficacy (exceeding a 1.5–2.0 order of magnitude drop) occurring in regimes characterized by low external drive (*ν ≤* 1.5, light green trajectories). (Right) Global parameter sweep heat map illustrating the spatial distribution of ΔLog Beta Power across relative inhibitory strength (*g*) and external drive ratio (*ν*) under standard high-frequency stimulation (130 Hz). Colored circular markers anchor the explicit single-point network coordinates mapped in the line profiles on the left. Coordinates located in the unexcited or silent regions along the bottom edge are explicitly masked in neutral gray.

The efficacy of beta-band suppression exhibits a strong state-dependence governed by the baseline network parameters. For networks with subthreshold external drive (*ν ≤* 1.0), significant attenuation of beta power can be achieved at relatively low stimulation frequencies, initiating at approximately 30 Hz. Crucially, this therapeutic effect saturates rapidly; beyond a 50 Hz threshold, further increases in the stimulation frequency yield diminishing returns, reaching a stable suppression plateau. Within intermediate network regimes characterized by low drive and low-to-moderate relative inhibition (1.0 *< g <* 2.0), a comparable sensitivity to lower pacing frequencies is observed. However, the absolute magnitude of beta power attenuation in this domain is substantially muted compared to sub-threshold networks. Here, the capacity for spectral suppression scales directly with the strength of local feedback inhibition, meaning networks with higher *g* values exhibit enhanced vulnerability to disruption. For these specific parameter configurations, increasing the stimulation frequency yields an exponential improvement in the efficacy of beta-band suppression.

Finally, for highly driven networks operating at elevated inhibitory weights (*g >* 2.0), low-frequency stimulation completely fails to alleviate pathological beta-band power. Specifically, in networks where *g ≈* 4.0, frequencies below a 100 Hz threshold demonstrate no reliable suppression of the endogenous rhythm. Beyond this critical frequency boundary, however, the intervention gains efficacy, with the residual beta power exhibiting an exponential decay as a function of increasing stimulation frequency, underscoring the necessity of high-frequency protocols in highly active, inhibition-dominated circuits.

#### 2.5.1 The Effect of Frequency Along a Trajectory in Parameter Space

To further show the dependence of stimulation effects at different frequencies at different regions of the *g, ν* parameter space, we looked at two linear trajectories (Figure 7).

**Figure 7:**
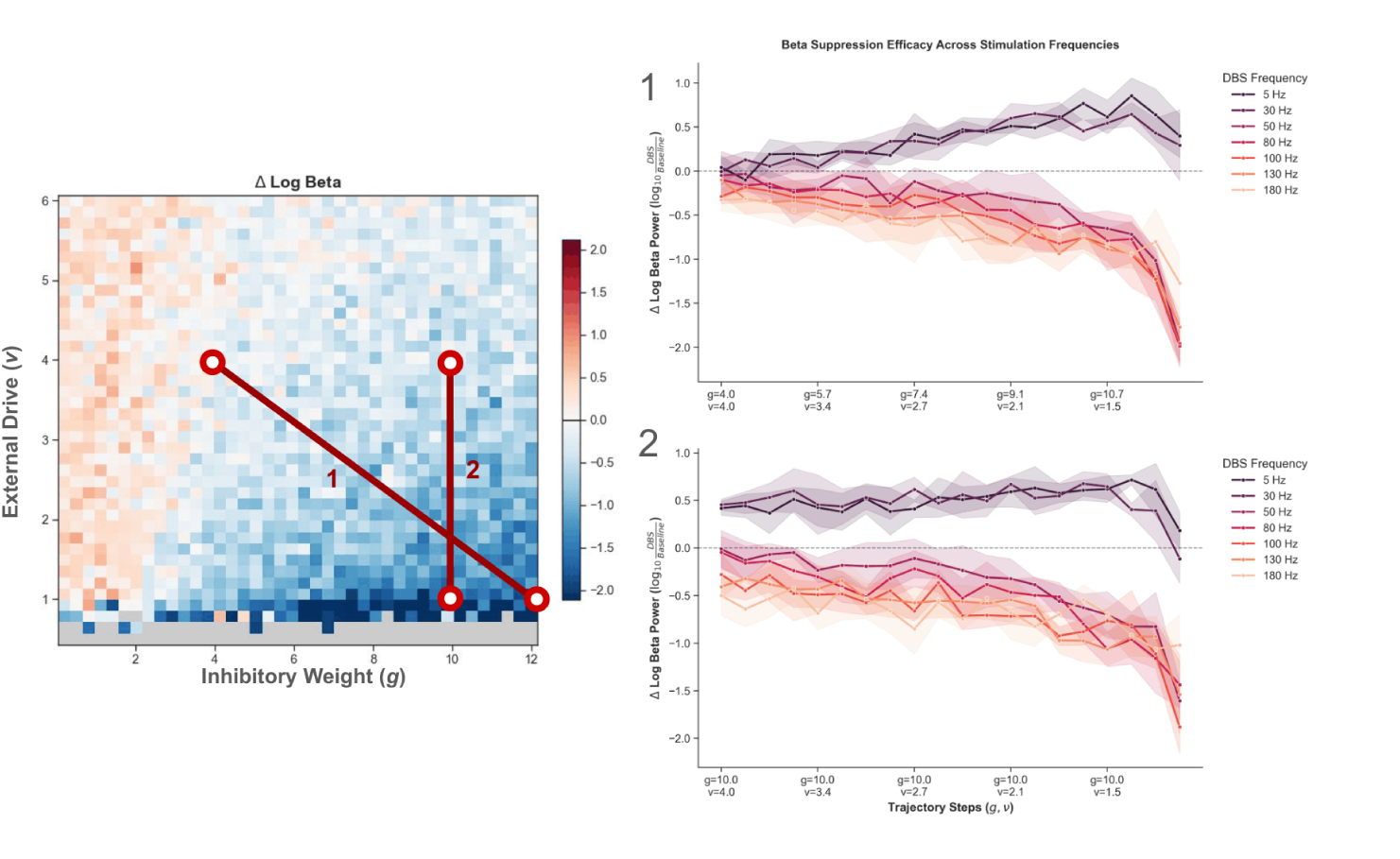
Context-dependent suppression of beta-band oscillations along parameter space trajectories. (Left) Spatial heatmap of the change in LFP log beta power (ΔLog Beta Power) under standard high-frequency stimulation (130 Hz) relative to baseline across the (*g, ν*) parameter landscape. Two strategic trajectories are highlighted to track stimulation efficacy across distinct state transitions: Trajectory 1 (diagonal path) transitions from a region of high external drive and weaker inhibition to a state of strong inhibition and low external drive; Trajectory 2 (vertical path) holds a strong inhibitory coupling constant (*g* = 10.0) while systematically lowering the external drive (*ν* : 4.0 *→* 1.0). Quiescent or silent network states along the bottom axis are marked in neutral gray. (Right) Suppression efficacy profiles plotted as a function of discrete trajectory steps under various DBS frequencies (5–180 Hz). Shaded bands represent *±*SEM. (1) Along Trajectory 1, high-frequency stimulation (*>* 100 Hz, warm orange/red curves) exhibits progressively stronger beta suppression as the network drops deeper into the low-drive, heavily inhibited regime. Low-frequency stimulation (5–30 Hz, purple curves) consistently amplifies beta oscillations across all steps. (2) Along Trajectory 2, therapeutic beta suppression is maximized when the external drive is minimized (*ν* = 1.5), matching the maximum desynchronization and suppression profiles observed in the overall state space.

The first trajectory follows a diagonal line from *g* = 4.0*, ν* = 4.0 to *g* = 12.0*, ν* = 1.0. As we had expected, we see at frequencies below 30 Hz, the stimulation has an amplifying effect on the beta-band activity of the network. The amplification effect gets progressively stronger along the trajectory as the network moves closer to the high inhibition, low external drive regime. There is a decrease in the amplification when the external drive becomes subthreshold; however the beta band activity still remains higher than in the baseline state. For frequencies greater than a 100 Hz, suppression of the beta-band activity is observed over the entire trajectory, with the effect increasing along the trajectory. The 130 Hz and 180 Hz stimulation does not show a great difference in their efficacy at suppressing beta oscillations, while the 100 Hz stimulation is less effective in networks with lower inhibition and higher external drive. The 50 Hz and 80 Hz stimulation are even less effective at lower inhibition and higher drive, and only starts showing an improvement in suppression efficacy towards the end of the trajectory. In the high inhibition, low external drive regime, the frequency of the stimulation has less of an effect as all stimulation frequencies of 50 Hz and greater producing similar suppression of beta-band activity.

We looked at a second trajectory that goes from a high inhibition, high drive state (*g* = 10.0*, ν* = 4.0) to a high inhibition, low drive state (*g* = 10.0*, ν* = 1.0). In networks with high inhibition, the level of external drive heavily influences the sensitivity to the frequency of stimulation. At high levels of external drive, the difference in effectiveness across stimulation frequencies is most evident: 180 Hz stimulation produces the greatest suppression, while 130 Hz and 100 Hz have a similar strength of effect. 50 Hz and 80 Hz do not reliably suppress beta-band activity in these networks. The 50 Hz stimulation does not have a reliable effect till the external drive is reduced to *ν ≈* 2.7. Across medium levels of external drive, higher frequencies consistently show a greater effect than lower frequencies. As the drive goes lower than *ν <* 2.0, the difference between the effects of the stimulation frequency start to shrink, with all frequencies of 50 Hz or greater producing a similar level of suppression as *ν →* 1.0. As previously observed, frequencies of 30 Hz or lower do not suppress and instead amplify the beta-band activity of the network, with the effect being greater in networks with lower drive.

## 3 Model and Methods

### 3.1 Neuronal Models

The neurons in the network were modelled as leaky integrate-and-fire neurons. The membrane potential of the neuron Vm changes in time according to the relationship

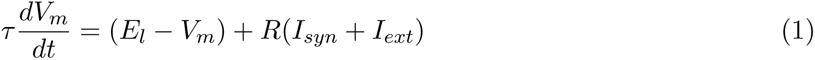

with the following constants: membrane time constant *τ*, leak reversal potential *E_l_*, membrane resistance *R*, and synaptic and external currents *I_syn_*, *I_ext_*.

When the membrane potential reaches a threshold value *V_thr_*, the neuron fires an action potential and membrane potential resets to *V_reset_*. After every action potential, the neuron has a refractory period *r_p_*.

Default values for the parameters are provided in Appendix A

To model the separate axonal and somatic responses to stimulation, we use a version of the Parrot neuron from the NEST neurosimulator. The Parrot neuron is a spike repeater that has no internal dynamics. It immediately fires an action potential when it receives a presynaptic input.

### 3.2 Synaptic Models

The synapses in the network are of two types: non-plastic exponential synapses and plastic Tsodyks-Markram depressing synapses [19]. The ratio of plastic to non-plastic synapses in the network is governed by the parameter *γ_plas_*.

The exponential synapses are governed by the equation

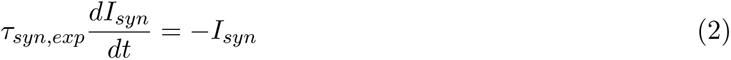

Parameter values for Tsodyks-Markram synapses are provided in Appendix B

### 3.3 Stimulation Model

The model of stimulation incorporates widespread presynaptic activation and axonal activation of the target neuron. When a stimulation pulse reaches a target neuron, it activates the Parrots of all the neurons presynaptically connected to it. This results in post-synaptic activation of the target neuron from all its afferent connections. The stimulation pulse also activates the Parrot of the target neuron, causing activation of all its efferent neurons. The activation of a Parrot neuron causes a post-synaptic activation of all the neurons it is connected to. As such, the activation of the target’s afferent Parrots causes activation of their other connections antidromically.

### 3.4 Network Model

The network used was a modification of the Brunel network comprising an excitatory and an inhibitory population with sparse connectivity between the populations, and recurrently to itself. The two populations also receive a common external input from a set of Poisson spike trains. The rate of the Poisson spike trains is governed by the parameter *ν*, and the relative weight of the inhibitory to excitatory connections is controlled by the parameter *g*. The network consisted of 1000 excitatory and 250 inhibitory neurons. The probability of connection was identical for excitatory-excitatory, excitatory-inhibitory, inhibitory-excitatory, and inhibitory-inhibitory connections and was set at 10%. Of all the connections, a certain fraction were plastic Tsodyks-Markram depressing synapses, while the rest were exponential synapses.

### 3.5 Simulations and Recording

Simulations were performed using a forward Euler method in Brian2 with a step size of 1 ms. For analysis, the first 100 ms were ignored. When stimulation was applied, the first 100 ms after stimulation onset were also ignored for calculating metrics. The membrane potential as well as the excitatory and inhibitory post-synaptic currents were recorded for the neurons. The analog for the Local Field Potential was calculated by the method used by [20] using the summed absolute postsynaptic currents.

### 3.6 Metrics

We used a diverse set of metrics to assess the changes in the network state. First we used the mean firing rate *S* that was calculated as follows:

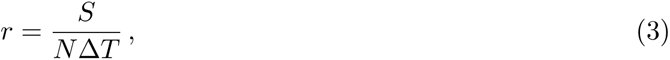

where *S* is the number of spikes in the time interval of length Δ*T* recorded from *N* neurons. Second, we used the Coefficient of Variation (*CV*) of the Inter-spike Intervals (ISIs) that was calculated as follows:

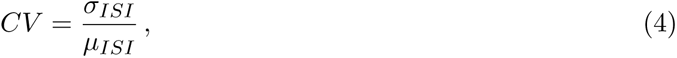

where *σ_ISI_* and *µ_ISI_*denote the standard deviation and mean of the ISIs.

Furthermore, we assess the level of synchrony in the network to characterize the network state. The Synchrony was measured as the pairwise-cross correlation of the spike trains. The spike trains were binned into intervals of width 2 ms. The cross-correlation matrix of the binned spike trains was calculated, and the mean of the upper triangular portion was denoted as the measure of synchrony. The beta power was calculated by computing the Power Spectral Density of the Local Field Potential using the Welch periodogram. The area under the curve in the beta frequency range (13-30 Hz) was numerically calculated by trapezoidal integration.

## 4 Discussion

### 4.1 Summary of Results

The primary objective of this work was to extend the classic Brunel network framework to include electrical neurostimulation, allowing for a systematic investigation of stimulation parameters across diverse network topologies and intrinsic activity regimes. By charting these interactions across the *g* and *ν* parameter space, we characterized the state-dependent effects of stimulation on collective dynamics, including population firing rates, coefficient of variation (CV), global synchrony, and pathological beta-band power suppression. We showed that high-frequency stimulation (HFS)—the clinical standard for therapeutic deep brain stimulation (DBS)—consistently disrupts pathological rhythms. Crucially, however, its secondary impact on other network characteristics is explicitly constrained by the network’s baseline state. Furthermore, we demonstrated that the minimum effective dosage required to achieve robust rhythm disruption is highly state-dependent, with therapeutic efficiency scaling monotonically with stimulation frequency.

### 4.2 Network Regime and State-Dependent Beta Suppression

A central challenge in computational neurostimulation is mapping abstract parameter topologies (such as the Brunel *g* and *ν* phase space) to specific pathological subcortical nuclei, such as the subthalamic nucleus (STN) or the globus pallidus internus (GPi) in Parkinson’s disease. While the absolute coordinates of these structures within an E-I balance grid remain difficult to define experimentally, our model offers valuable predictive insights based on relative network dynamics. As the network shifts away from a tightly balanced E-I state toward a more heavily inhibited regime (increasing *g*), we observed a pronounced increase in beta-band suppression. Physiologically, this highly inhibited regime aligns closely with the baseline environment of the GPi, which receives dense, high-frequency GABAergic inputs from the striatum and globus pallidus externus (GPe) [21]. Experimentally, the transition from a healthy to a Parkinsonian state in the GPi is characterized by a transition from tonic irregular firing (typically 60–80 Hz) to faster, more regular firing (*>* 100 Hz) [22]. When we look at our model’s response within these highly inhibited, high-baseline drive zones, we find that robust beta suppression is achieved only at traditional clinical frequencies (*∼* 130 Hz) with lower frequency stimulation failing to produce reliable results. Conversely, shifting to the left of the parameter space, where excitation and inhibition is more balanced and the external drive is high, activity is similar to the ventral intermediate nucleus (VIM) of the thalamus in essential tremor pathology, characterised by tonic firing along with a subset of neurons exhibiting synchronized population bursts. Stimulation in this case elicits transient phase-locking followed by an un-oscillatory, steady-state constant firing pattern, mirroring the clinical efficacy of VIM-DBS in treating essential tremor [2, 23]. Finally, we observe that networks with high inhibition and low drive that show low baseline firing are most susceptible to low-frequency stimulation. This may be compared to stimulation of the pedunculopontine nucleus [24, 25], where frequencies below 50 Hz have been shown to reduce motor symptoms in select Parkinsonian patients [26]. These findings suggest that the clinical target’s intrinsic E-I balance governs its susceptibility to stimulation. Subcortical targets with more dominant inhibitory configurations and lower external drive are inherently more susceptible to lower-frequency disruptive inputs, whereas structures operating under lower baseline inhibition require higher frequency stimulation to disrupt synchronized oscillations.

### 4.3 Model Scope, Circuit Motifs, and Limitations

While our model successfully captures the broad, parameter-dependent properties of neurostimulation, it is important to recognize the limitations of an isotropic, randomly connected network model. The real basal ganglia-thalamocortical loop relies on distinct topological motifs, such as closed-loop feedback, feedforward inhibition, and varied synaptic dynamics, which are simplified within this framework. Consequently, certain network responses may be influenced by complex architectural features that a standard Brunel model cannot fully replicate. Nevertheless, we anticipate that the fundamental relationships identified here, specifically the interaction between E-I balance, external drive, and structural susceptibility to external periodic stimulation, will remain preserved across more complex architectures. This model serves as a necessary foundation for uncovering the core mechanisms of network-level DBS-modulated collective dynamics before introducing more intricate anatomical complexities.

### 4.4 Criticality and Homoeostasis

Assessing these shifts through the lens of neural criticality provides a valuable framework for understanding network stability. Evidence indicates that Parkinsonian beta oscillations represent an over-synchronized, subcritical state that restricts the network’s capacity for information routing [27]. Effective neurostimulation appears to disrupt this pathological state, shifting the network’s dynamics back toward a more balanced, critical regime [28].

However, this transition must be interpreted conservatively. Rather than driving the network precisely to a critical point, stimulation likely acts as a stabilizing force that prevents the system from getting stuck in highly synchronized, rigid states. This shift broadens the network’s dynamic range and restores functional variability, which correlates directly with clinical symptom alleviation.

### 4.5 Methodological Framework and Extension

Beyond the specific insights into Parkinson’s disease dynamics, the Brunel network provides a modular framework for testing other neurostimulation modalities, with modifications that can be made to best reproduce specific stimulation features. This framework can easily be extended to model global non-invasive interventions, such as Transcranial Magnetic Stimulation (TMS).

Integrating a macro-scale model of TMS-induced electric fields into this network would allow one to investigate how global, low-density current distributions interact with underlying E-I dynamics [29, 30]. This approach would make it possible to directly compare the network-level mechanisms of focal, high-density invasive stimulation (DBS) with widespread, non-invasive cortical stimulation (TMS) within the exact same computational framework. Such comparative modeling is an essential step toward designing multi-scale, patient-specific stimulation protocols that optimize therapeutic outcomes across various neurological disorders.

## A List of Parameters and their Default Values

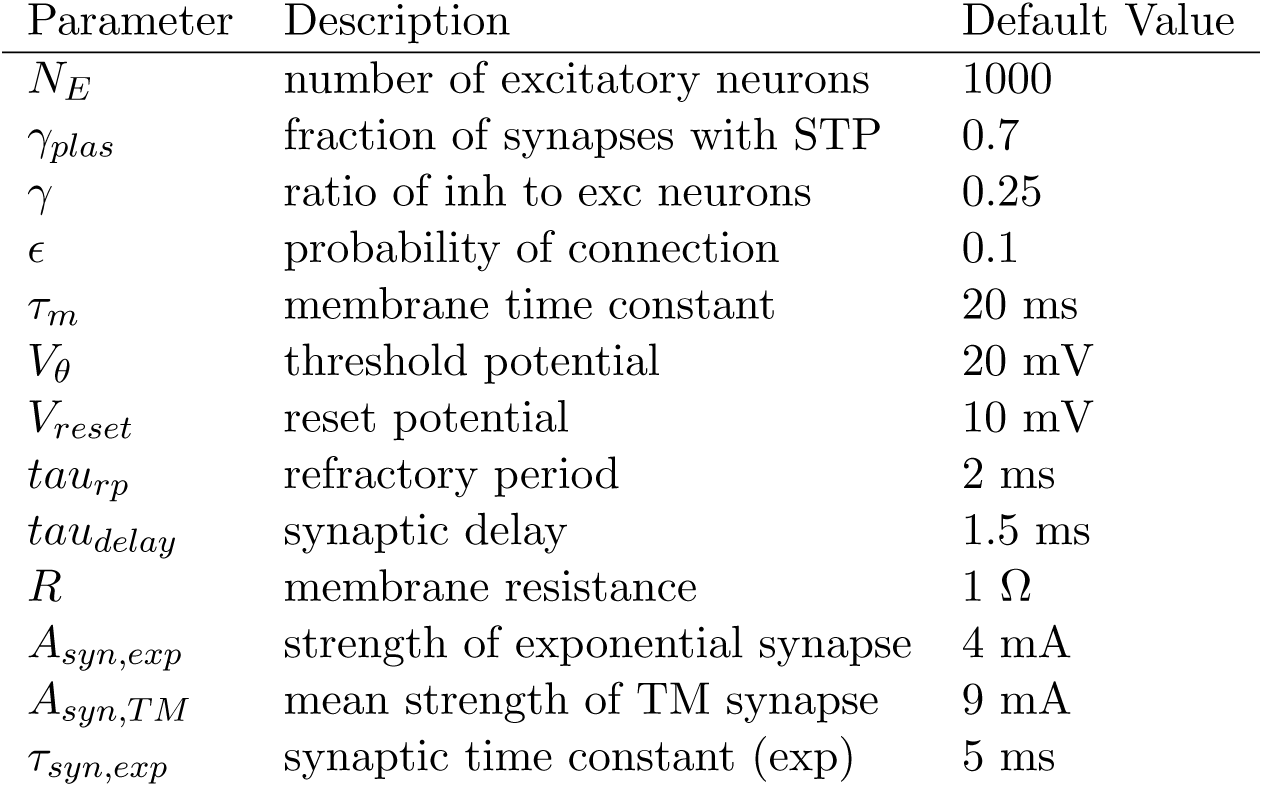

| Parameter | Description | Default Value |
| --- | --- | --- |
| $N_E$ | number of excitatory neurons | 1000 |
| $\gamma_{plas}$ | fraction of synapses with STP | 0.7 |
| $\gamma$ | ratio of inh to exc neurons | 0.25 |
| $\epsilon$ | probability of connection | 0.1 |
| $\tau_m$ | membrane time constant | 20 ms |
| $V_\theta$ | threshold potential | 20 mV |
| $V_{reset}$ | reset potential | 10 mV |
| $\tau_{rp}$ | refractory period | 2 ms |
| $\tau_{delay}$ | synaptic delay | 1.5 ms |
| $R$ | membrane resistance | 1 $\Omega$ |
| $A_{syn,exp}$ | strength of exponential synapse | 4 mA |
| $A_{syn,TM}$ | mean strength of TM synapse | 9 mA |
| $\tau_{syn,exp}$ | synaptic time constant (exp) | 5 ms |

## B Tsodyks-Markram Parameters

| Parameter | Description | Exc Value | Inh Value |
| --- | --- | --- | --- |
| $U$ | probability of release | 0.7 | 0.7 |
| $\tau_{rec}$ | recovery time constant | 300 ms | 850 ms |
| $\tau_i$ | inactivation time constant | 5 ms | 2 ms |

## Notes

### Competing Interest Statement

The authors have declared no competing interest.

### Funding

- Natural Sciences and Engineering Research Council of Canada
- Canadian Institutes of Health Research
- Max Planck-University of Toronto Centre (MPUTC)

